# Overexpression of the *Arabidopsis* thaumatin-like protein 1 in transgenic potato plants enhances resistance against early and late blights

**DOI:** 10.1101/621649

**Authors:** Gul Shad Ali, X Hu, A.S.N. Reddy

## Abstract

The *Arabidopsis* thaumatin-like protein 1 (ATLP1), which belongs to the PR5 family of pathogenesis-related proteins, is induced by pathogen attacks and by systemic acquired resistance (SAR)-inducing compounds. To test whether the overexpression of ATLP1 will enhance fungal resistance in transgenic plants, the cDNA of this gene under the control of a constitutive promoter was transferred into *Solanum tuberosum* Cv. Desiree. The expression of the introduced gene was confirmed by Northern and Western blot analyses. Western blot analysis performed with intercellular fluid (ICF) showed that ATLP1 is secreted to apoplast. Several independent transgenic lines with high-level expression of the ATLP1 were assayed for resistance against early blight (*Alternaria solani*) and late blight (*Phytophthora infestans*). The rate of *Alternaria* lesions development was significantly reduced in the ATLP1 transgenic lines as compared to a control line. Percent reductions in area under the disease progress curve (AUDPC) values for the ATLP1 transgenic lines as compared to control line ranged between 52 and 82%. In response to *P*. *infestans*, infection efficiency (IE), lesion size (LS) and sporulation capacity (SC) were significantly reduced in ATLP1 transgenic lines as compared to a control line. On the average, IE, LS and SC were reduced by 18, 22 and 20%, respectively, for all transgenic lines with a maximum reduction of 25% in IE, 25% in LS and 32% in SC. Resistance assays against *P*. *infestans* using whole plants showed 40-70% reduction in symptoms as compared to control. These results suggest that constitutive expression of a heterologous ATLP in potato confers enhanced resistance against early and late blights.

## Introduction

In response to pathogen attacks, plants induce a complex of defense mechanisms, which in most cases involves localized programmed cell death (PCD) commonly called hypersensitive response (HR) (Staskawicz et al., 1995; Baker et al., 1997). HR not only results in confining the invading pathogen to the area of infection but also triggers non specific defense responses throughout the plant known as systemic acquired resistance (SAR) (Ryals et al., 1996). Concomitant with the induction of HR and SAR a plethora of defense-related genes are expressed (Ryals et al., 1996; Fritig et al., 1998). These genes are involved in strengthening cell walls by covalently cross-linking cell-wall matrix polymers (Bowles, 1990), in synthesizing antimicrobial compounds, such as phytoalexins (Dixson, 1986; Guo and Paiva, 1995), and in coding pathogenesis related proteins, some of which encode lytic enzymes, e.g. chitinases and glucanases (Lamb et al., 1989) and other antimicrobial proteins (Bowles, 1990). In the majority of plant disease resistance responses most of these defense-related genes are activated, suggesting that a common resistance activation mechanism is prevalent in most plants (Kunkel, 1996). However, despite the presence of such an elaborate defense system many devastating plant diseases do occur suggesting that either these defense mechanisms are too weak or appear too late to be effective for protecting the invaded plant (Fritig et al., 1998). Therefore, the constitutive expression of these defense-related genes in transgenic plants has been suggested to enhance the resistance of these plants to pathogenic organisms (Lamb et al., 1992).

Perhaps the most intensely studied SAR genes are the ones encoding the so-called pathogenesis-related (PR) proteins, originally discovered as novel proteins accumulating in tobacco after TMV infection (Van loon and Van Kammen, 1970; Van Loon, 1985), and subsequently found to be associated with an array of plant-pathogen interactions (Linthorst, 1991; Ryals et al., 1996; Bliffeld et al., 1999). Several of the PR-proteins, e.g., PR-1a, PR-2 (glucanases), PR-3 (chitinases), PR-4 and PR-5 (osmotin, thaumatin-like proteins) are involved in resistance to the primary pathogen (Bowles, 1990) as well as subsequent pathogens (McIntyre et al., 1981) and exhibit antimicrobial activity *in vitro* (Collinge et al., 1993; Hammond-Kosak and Jones, 1996). Furthermore, evidence for the role of these proteins in defense against pathogens has been obtained by showing that overexpression of some of them alone or in combinations in transgenic plants resulted in enhanced protection against fungi (Broglie et al., 1991; Alexander et al., 1993; Hain et al., 1993; Vierheilig et al., 1993; Liu et al., 1994; Zhu et al., 1994; Jach et al., 1995; Grison et al., 1996; Punja and Raharjo, 1996; Bliffeld et al., 1999; Chen et al., 1999; Datta et al., 1999; Yu et al., 1999).

Potato crop sustains significant yield losses due to fungal diseases (Hooker, 1981). The most damaging fungal diseases are early and late blights caused by *Alternaria solani* and *Phytophthora infestans*, respectively. Crop losses can go up to 100 percent. In developing countries late blight alone is costing farmers an estimated 2.75 billion dollars each year (Annonymous, 2005). Likewise in the Columbian basin about 22.3 million dollars worth of fungicides were sprayed for the control of late blight in 1998 alone (Johnson et al., 2000). Fungicide application is currently the best control strategy for controlling fungal diseases in potato fields. However, due to the environmental concerns of fungicides, and the appearance of fungicide resistant strains of fungal pathogens, particularly of *P*. *infestans*, development and deployment of resistant cultivars is inevitable. Using traditional breeding techniques, a number of useful resistance genes have been introduced into commercial cultivars. However, such resistance has met with very limited success, first, because it is difficult to breed because the sources of resistance are often derived from wild species and second, it is short-lived and difficult to maintain for long periods because of the rapidly evolving virulent pathogen populations. Under these circumstances genetic engineering allows for the rapid development of resistant cultivars by introducing one or a few genes for resistance from heterologous sources into commercially acceptable varieties. The constitutive expression of a variety of antifungal genes in transgenic plants has been shown to have enhanced resistance against fungal pathogens both under greenhouse and field conditions. For potato, enhanced resistance against fungal pathogens by the introduction of genes for osmotin-like protein (Liu et al., 1994; Zhu et al., 1996; Li et al., 1999) glucose oxidase (Wu et al., 1995), the *Trichoderma* endochitinase (Lorito et al., 1998) and utilizing novel protection mechanisms (Strittmatter et al., 1995; Yu et al., 1999) have been reported.

The Arabidopsis thaumatin-like protein 1 (ATLP1) is a 25 kD cysteine-rich protein that is induced in response to pathogen attacks and to systemic acquired resistance (SAR) compounds e.g. salicylic acid (SA) (Hu and Reddy, 1997). This protein has the characteristic 16 cysteine residues interlinked through 8 disulfide bonds commonly found in the members of PR5 family of pathogenesis related proteins. *In vitro* antifungal assays performed with a purified closely related counterpart has demonstrated inhibitory effects against *Verticillium alb*-*atrum*, *V*. *dahliae*, *A*. *solani*, *Fusarium oxysporium* and *Trichoderma reesei* (Hu and Reddy, 1997). In addition, the constitutive expression of a rice thaumatin-like protein in rice (Datta et al., 1999) and wheat (Chen et al., 1999), and osmotin, a closely related PR5 protein, in transgenic potato plants (Liu et al., 1994; Zhu et al., 1996) lead to enhanced resistance against fungal pathogens. These findings suggest that ATLP1 functions as a defensive gene and could enhance resistance when overexpressed in transgenic potato plants. The objectives of this study were to produce transgenic potato lines overexpressing ATLP1, and to evaluate the resistance of ATLP1-transgenic potato plants against *P*. *infestans* and *A*. *solani*.

## MATERIALS AND METHODS

### Vector contructs and plant transformation and regeneration

A cDNA fragment consisting of the entire open reading frame, and 5’ and 3’ untranslated regions of Arabidopsis thaumatin-like protein 1 (ATLP1) was cloned in the pBluescript. The resultant plasmid was digested with S*ma*I/X*ho*I and the released 1008 bp fragment was cloned in the S*ca*I/X*ho*I sites between the CaMV 35S promoter and terminator of the plant transformation binary vector pGA748. To generate antisense construct, the cDNA of ATLP1 in pBluescript was excised with X*ho*I/B*am*HI and inserted in the X*ho*I/B*gl*II sites of the pGA748. The chimeric sense and antisense gene constructs, designated ATLP1-S and ATLP1-As respectively, were introduced into *Agrobacterium tumefaciens* strain LBA 4404 by electrophoration. Potato Cv. Desiree was transformed with ATLP1-S and ATLP1-As by *Agrobacterium*-mediated leaf disk transformation method. Briefly, leaf disc were incubated with the *Agrobacterium tumefaciens* (OD_600_=0.1) strain LBA 4404 carrying ATL1-S or ATLP1-As vectors for 10 minutes and transferred to callus-induction medium consisting of Murashige and Skoog (MS) salts, glucose 16 g L^−1^, zeatin ribose, 1 mg L^−1^, NAA, 20 *μ*g L^−1^, GA3 20 *μ*g L^−1^, AgNO_3_ 10 mg L^−1^, carbenicillin 100 mg L^−1^, kanamycin100 mg L^−1^ and phytagar 8 g L^−1^, pH 5.7. After incubation for two days in the dark at 24 °C, leaf disks were transferred to fresh plate of the same medium every 10 days till calli and shoots developed. Well-developed shoots were transferred to root-induction medium which consisted of half-strength MS salts, sucrose 20 g L^−1^, carbenicillin 100 mg L^−1^, kanamycin 100 mg L^−1^ and phytagar 8 g L^−1^. After confirmation by Northern blot analyses, independently transformed lines were multiplied on the same selection medium and transferred to soil (Metro-Mix 350, American Clay works & Supplies, Denver, CO) in a green house maintained at 23±2 °C and 16:8 light:dark cycle.

### RNA blot analyses

Total RNA was separated on 1% Agarose gel and blotted onto Nylon membranes. Equal loading was verified by ethidium bromide staining of the gel. The RNA gel blots were hybridized to a ^32^P-labelled ATLP1-specific probe and exposed to X-ray film per standard procedures.

### Production of antibody against ATLP1 and immunoblot analyses

For antibody production, the ATLP1 cDNA cloned in pET28a was expressed in *E. coli* BL21 strain as described previously (Hu and Reddy, 1997). Total protein from induced cells was resuspended in Laemalli buffer and run on a 14% SDS-polyacrylamide gel. The band corresponding to the size of ATLP1 was excised from the gel, eluted and used for raising antibody in rabbit at the Macromolecular Resources Facility at Colorado State University, Fort Collins, CO, USA. Antisera collected at various time-points were verified to react with the bacterially expressed ATLP1 using western blot analyses.

Total protein from leaf tissues, macerated in liquid nitrogen, was extracted with Tris-buffer (Tris-HCl 50 mM, EDTA 5 mM, Sucrose 250 mM, DTT 5 mM, pH 7.5) by centrifugation at 5000 g at 4 °C. Two volumes of the crude extract was mixed with one volume of 3x Laemalli gel loading buffer and heated at 90 °C for 5 minutes. After centrifugation for 5 minutes, 30 *μ*L of the total protein was separated on 12% SDS-containing polyacrylamide gel and transferred to nitrocellulose membrane using Bio-Rad Trans-Blot electrophoretic transfer system. Membranes were blocked with 3% gelatin in TBS overnight at 30 °C. The blots were probed with antiserum (1:2000 dilution) raised against gel-purified ATLP1 using alkaline phosphatase(AP)-conjugated goat anti-rabbit IgG (1:10000 dilution) as the secondary antibody and 5-bromo,4-chloro,3-indolylphosphate(BCIP)/nitroblue tetrazolium (NBT)(Sigma Chemical Co. MS) as the substrates (Safadi et al., 2000).

### Infection assays

Resistance of transgenic lines to early and late blights were evaluated by both detached leaf and whole plant assays in a growth chamber. *A*. *solani* isolated from an infected field-grown potato plant was cultured on half-strength potato dextrose agar at room temperature (approximately 25°C) under white fluorescent light on a laboratory bench for 10 days. Sporulation was induced by rubbing fungal mycelia on the plates and left opened in inverted position for 36 hours. Spores were collected in half-strength potato dextrose and the concentration adjusted to 10^5^ spores mL^−1^. For detached leaf assays, eight leaflets of equal age and size from fully expanded leaves, approximately the third from the top of the plants were excise and placed adaxial side up on water agar contained in petri dishes. Each leaflet was inoculated with 10 *μ*L spore suspension on one location and placed at room temperature under white fluorescent light. After development of symptoms, inoculated leaves were scanned using the Leaf Lumina scanning system (Leaf System, Southboro, Massachusett) on the 2^nd^, 3^rd^, 4^th^ and 5^th^ day after inoculation. Disease leaf area was determined on the scanned photos using the histogram function under the Image menu of Adobe PhotoShop software (Adobe Systems Inc. CA). Areas under the disease progress curve (AUDPC) were calculated from the lesion areas recorded in pixels and analyzed using the GLM procedure of the SAS software (SAS Institute Inc., Carey NC). Differences between each transgenic line and the control, ATLP1-As8, were tested by the Students’ t-test using the solution option in the model statement of the GLM procedure. Also percent reduction in AUDPC of transgenic lines as compared to control was calculated from the averaged values. Total chlorophyll content in potato leaves was measured as described (Arnon et al., 1974).

*P*. *infestans* strain US8 was cultured on detached leaves of potato Cv. Desiree placed on water-agar plates at 16° C for 10 to 14 days. Sporangia were collected in water. The sporangial suspension was adjusted to 10^5^ sporangia mL^−1^ and incubated at 4 °C for 4 hours to induce zoospores release. Each leaflet was then inoculated with a 10 *μ*L zoospore suspension at one location and placed in growth chamber maintained at 17°C, 100% RH and 16:8 hr light:dark regime. Five days post inoculation leaves were scored for disease response on a scale of 0-4 with 0= no infection/non sporulating lesions, 1= 1-25%, 2= 26-50%, 3= 51-75% and 4= 76-100% of the lesion covered with spores. Percent infection efficiency (IE), defined as the percent of inoculation sites showing sporulating lesions, of the transgenic lines over ATLP1-As8 were calculated. Sporulation capacity (SC) was measured as sporangia per square centimeter of the lesion. IE, disease area and sporulation capacity were analyzed by the GLM procedure of SAS statistical package (SAS®, SAS Institute Inc., Carey NC) with the solution option for mean comparisons. Before analysis data were checked for normality.

For whole plant assays, transgenic and control potato plants were generated from tissue culture plants as mentioned above. Five to eight seven-week old plants grown in 9-cm pots were inoculated with *P*. *infestans*-sporangial suspension prepared as above. Immediately after inoculation, plants were placed in the dark for the first 24 hours in a growth chamber maintained at 17°C. This was to improve the infection of *P*. *infestans*. Afterward, plants were placed in the same growth chamber under a 16:8 light:dark regime. Ten days after inoculation, percent leaf area showing disease symptoms were determined. Percent disease-leaf area were log transformed and analyzed by the GLM procedure of SAS software. Mean disease areas of ATLP1-transgenic lines were compared to control by students’ t-test. The experiment was repeated once.

## RESULTS

### Transformation and regeneration of potato

Figure 1A shows the gene constructs used to generate transgenic potato plants used in this study. The plasmid ATLP1-S and ATLP1-As were used to transform potato Cv. Desiree by *Agrobacterium*-mediated leaf disk transformation method. A total of 53 independent lines were generated, selected and maintained on kanamycin-containing medium as described in materials and methods.

**Figure 1.**
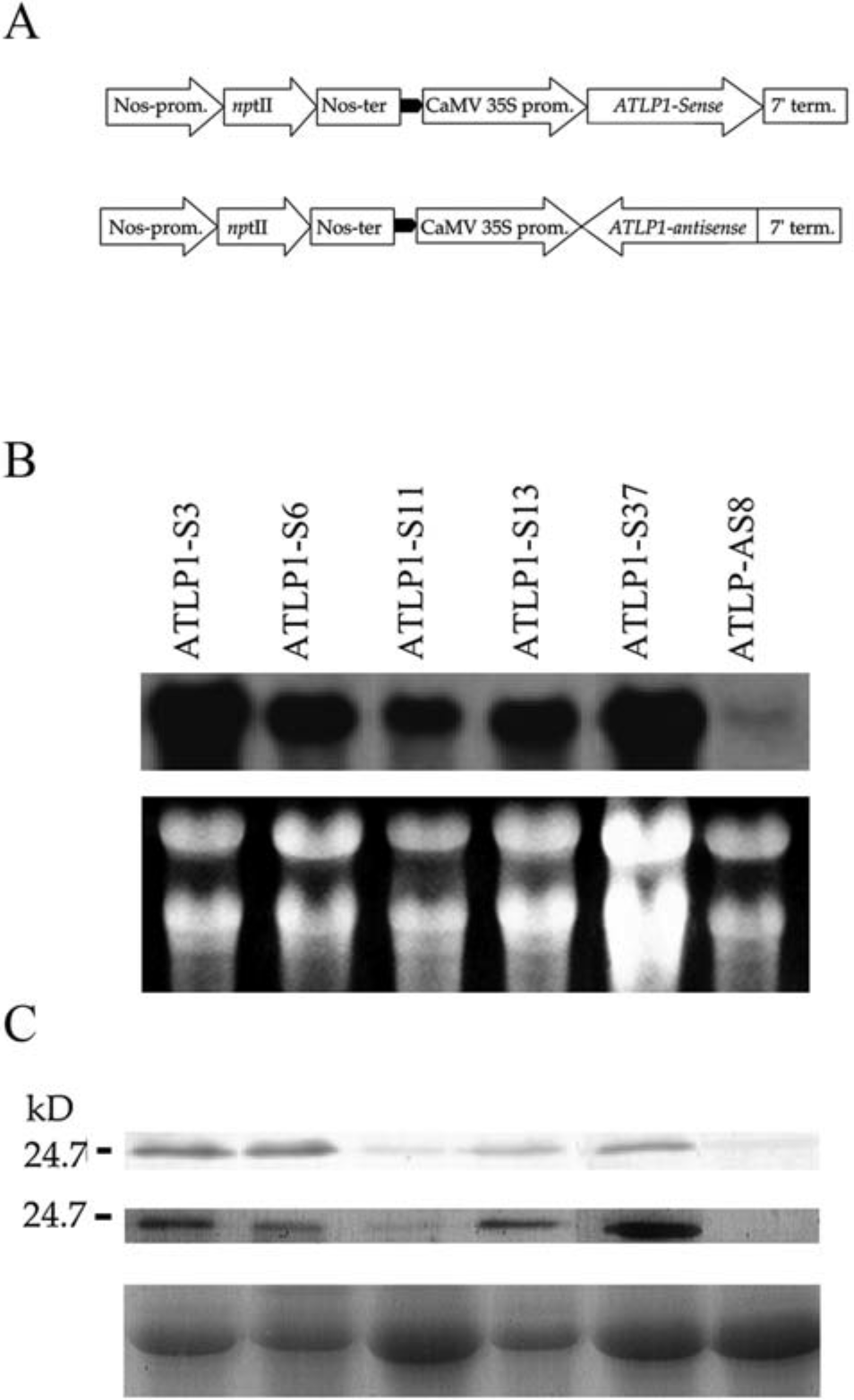
Molecular analyses of the ATLP1 transgenic potato plants. **A)** Gene fusions for potato transformation. Full length cDNA of Arabidopsis thaumatin-like protein was fused to the Cauliflower Mosaic Virus 35S promoter in sense orientation, ATLP1-Sense and antisense orientation, ATLP1-antisense. *Npt*II = the neomycin phophotransferase II gene, Nos-prom. = Nopaline synthase-Promoter, 7’ term. = polyadenylation signalof the *Agrobacterium tumefaciens* T-DNA gene 7. **B**) Analysis of ATLP1 mRNA production. Equal amounts of total RNA were run on a 1% gel, transferred to nylone membrane and hybridized with a ^32^P-labelled probe speceific for ATLP1. Lower panel is an ethidium-bromide stained gel showing amount of total RNA loaded per well. ATLP1-S3 to ATLP1-S37 are sense transformants; ATLP1-As8 is an antisense transformant. **C)** Analyses of ATLP1 protein production in the same set of plants as given in B. Total leaf protein (upper panel) or intercellular fluid (middle panel) of ATLP1-transgenic and control plants were subjected to SDS-PAGE and detected with antibody raised against ATLP1. The bottom panel is a coomassie stained protein gel showing the amount of protein loaded per lane.

### Molecular analyses of transgenic plants

Of the kanamycin resistant primary transformants, 5 sense lines were screened for the level of ATLP1 by Northern and Western blot analyses. Using an ATLP1-specific probe, several sense transgenic lines had high mRNA levels while antisense transgenic line ATLP1-As8 had no or very low detectable transcription (Figure 1B). Also evident from Figure 1B is the substantial variation in the level of ATLP1 transcripts among the transformants. Protein expression of ATLP1-S lines that had been shown to express ATLP1 was determined by immunoblot analysis using a polyclonal antibody raised against ATLP1 protein. Figure 1C illustrates the expression levels of ATLP1 in total leaf protein extractions. Again, like mRNA levels, protein expression was considerably high in all ATLP1 sense lines but not in ATLP1-As control line (Figure 1C, upper panel). This suggests that the anti-ATLP1 antibody did not cross react with the potato thaumatin-like proteins and that the observed protein bands are from the expression of the ATLP1 and not the endogenous potato PR5-like proteins. Since the chimeric ATLP1 gene introduced into potato plants had its own 20 amino acids secretory signal peptide, the secretion and localization of the protein was examined by Western blot analysis performed with intercellular fluid (ICF) from these transgenic lines. A specific band of 24.0 kD was detected in the intercellular fluids of ATLP1-sense transgenic lines but not in ATLP1-antisense or non-transformed control lines (Figure 1C, middle pannel).

### Early and late blight resistance assays

Transgenic potato plants expressing sense ATLP1 were compared to an antisense ATLP1 transgenic line (ATLP1-As8) for disease resistance response to inoculation with *A*. *solani* and *P*. *infestans* in detached leaf and whole plant assays. To quantify resistance response to *A*. *solani*, necrotic lesions, which appeared two days after inoculation, were measured from 2 to 5 days post-inoculation (dpi). Disease development was rated as area under the disease progress curve (AUDPC) based on expanding lesion sizes. Pairwise t-tests of ATLP1-S lines with antisense ATLP1-As8 performed on the combined data of 2 independent experiments showed significant differences (p=0.01 - 0.0001) for all pairs of comparison (Table 1). This is illustrated in Figure 2A where progression of lesion development on leaves from representative ATLP1-transgenic lines and ATLP1-As8 control inoculated with *A*. *solani* are shown. The lesions expansion data on ATLP1-transgenic lines are quantitatively presented as AUDPC in figure 2B. The rates of disease development on ATLP1-transgenic lines were reduced by 52-82% when compared to ATLP1-As8 (Figure 2B and Table 1). *Alternaria* spp are necrotrophic fungi that kill cells. Toxins produced by the fungus spreads outside the actual site of infection, which leads to the loss of chlorophyll from leaves, commonly termed as chlorosis (Hooker, 1981). Chlorosis caused by *A*. *solani* was quantified indirectly by measuring the total chlorophyll content of inoculated leaves in the ATLP1 transgenic and the control on the 5^th^ day after inoculation. In the ATLP1-As8 control line the chlorophyll level was only 21% compared to that of its corresponding un-inoculated control (Figure 2C). Whereas the ATLP1 transgenic lines had 65 to 82% of their normal chlorophyll content indicating that the overexpression of ATLP1 has impaired the virulence of *A*. *solani* (Figure 2C).

**Table 1.** Resistance of ATLP1-trasgenic potato plants to *A*. *solani*

| **Transgenic** | **% reduction over** |  |
| --- | --- | --- |
| **Lines** | **control** | **p-value<sup>a</sup>** |
| ATLP1-S3 | 81.9 | 0.0109 |
| ATLP1-S6 | 80.2 | 0.0001 |
| ATLP1-S11 | 53.2 | 0.0001 |
| ATLP1-S13 | 74.0 | 0.0001 |
| ATLP1-S37 | 52.7 | 0.0001 |
| ATLP1-As8 | 00.0 | - |

<sup>a</sup> The differences between ATLP1-S transgenic lines and a control ATLP1-As8 line was tested by student's T-test. P-values are probabilities of getting a larger value of T if the difference between an ATLP1-S transgenic and ATLP1-As8 control line was truly equal to zero.

**Figure 2.**
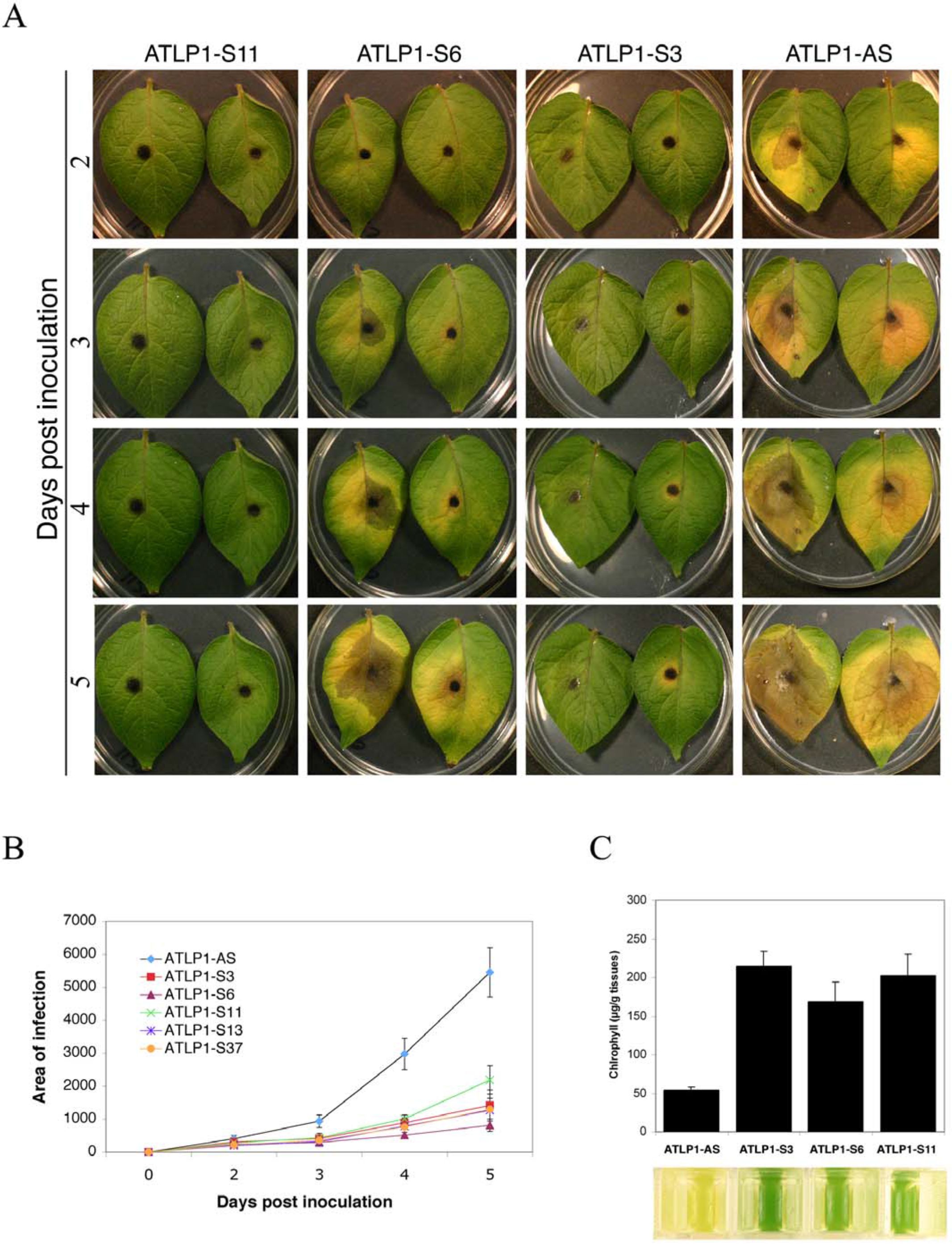
Response of representative ATLP1-transformed potato plants and a control to early blight (*A*. *solani*) in a detached-leaf assay. **A)** Leaflets were inoculated with a spore suspension of *A*. *solani* and placed under fluorescent light at RT. Each panel consists of two leaves from a representative line. Black spots in the middle of lesions are inoculation points. **B)** Reduced rate of lesion development on the detached leaves of ATLP1-transformed potato plants to early blight (*A*. *solani*). On the 2^nd^, 3^rd^, 4^th^ and 5^th^ day after inoculation, leaflets were scanned and the digital photos used for measuring disease area. Each data point is the average of two independent experiments each consisting of at least 8 inoculation points. Vertical bars are standard errors. **C**) Chlrophyll content of the *Alternaria*-inoculated leaves. Chlorophyll contents were determined on the 5^th^ day after inoculation. Data given are mean ± SE of three replicates. Lower panel shows representative samples of the leaf extracts used in measuring the chlorophyll contents presented in the bar graph above.

Response to inoculation with *P*. *infestans* was assessed both in detached leaf and whole plant assays. In the detached leaf assays resistance was evaluated by measuring infection efficiency (IE), lesion size (LS) and sporulation capacity (SC) 5 or 6 days post-inoculation. These parameters were significantly reduced in all tested ATLP1-transgenic lines as compared to ATLP1-As8 control indicating that expression of ATLP1 affects several stages in the life cycle of *P*. *infestans* (Table 2). On the average IE, LS and SC were reduced by 18, 22 and 20% respectively for all transgenic lines. When all these three components were averaged, a 4-29% reduction was observed for the transgenic lines, with ATLP1-S3 with the highest rank (24%), followed by ATLP1-S6 (18%), ATLP1-S37 (16%), ATLP1-S11(13%) and ATLP1-S13 (4%). A range of sporulation classes, 0-4 with 0 being the lowest and 4 being the highest, were observed in the ATLP1 transgenic lines. In general, lesser sporulating lesions occurred more frequently in ATLP1-S lines than in ATLP1-As8 line (Table 3). Representative pictures from leaves inoculated with *P*. *infestans* are shown in figure 3A. Whitish gray areas in the middle of the leaves are spores of *P*. *infestans*.

**Table 2.** Infection efficiency (IE), lesion size (LS) and sporulation capacity (SC) of *P*. *infestans* on ATLP1-transgenic potato plants

| Lines | % of |  | % of |  | % of |  |
| --- | --- | --- | --- | --- | --- | --- |
|  | IE<sup>a</sup> | control | LS<sup>a</sup> | control | SC<sup>a</sup> | control |
| ATLP1-S3 | 63.9±0.01<sup>b</sup> | 76.7 | 1.16±0.08 | 74.9 | 4.9±0.8 | 78.2 |
| ATLP1-S6 | 62.5±9.7 | 75.0 | 1.20±0.09 | 77.5 | 5.8±0.9 | 92.5 |
| ATLP1-S11 | 75.0±9.6 | 90.0 | 1.23±0.07 | 79.4 | 5.8±0.5 | 92.3 |
| ATLP1-S13 | 75.0±13.9 | 90.0 | 1.58±0.08 | 100.0 | 6.2±1.3 | 98.3 |
| ATLP1-S37 | 83.4±8.4 | 100.0 | 1.28±0.08 | 82.6 | 4.3±0.4 | 67.8 |
| ATLP1-As8 | 83.4±8.4 | 100.0 | 1.55±0.10 | 100.0 | 6.3±1.0 | 100.0 |

<sup>a</sup> Infection efficiencies (IE) are the percent successful infection of all inoculation points;
Lesion sizes (LS) are in cm<sup>2</sup>; Sporulation capacities (SC) are given as spores per cm<sup>2</sup> X 10<sup>4</sup>
<sup>b</sup> Values after ± are standard errors
ATLP1-S3 to S37 are sense transgenic lines whereas, ATLP1-As8 is a control line

**Table 3.** Resistance of ATLP1-transgenic potato plants to *P*. *infestans*

| Lines | Frequency (%) of inoculated spots<br>with a sporulation class<sup>a</sup> of: |  |  |  |  | Total<br>number of<br>inoculated<br>spots |
| --- | --- | --- | --- | --- | --- | --- |
|  | 0 | 1 | 2 | 3 | 4 |  |
| ATLP1-S3 | 39.7 | 41.2 | 8.8 | 8.8 | 1.5 | 68 |
| ATLP1-S6 | 25.7 | 17.1 | 12.9 | 22.9 | 21.4 | 70 |
| ATLP1-S11 | 25.4 | 46.5 | 16.9 | 11.3 | 0.0 | 71 |
| ATLP1-S13 | 37.1 | 27.1 | 21.4 | 11.4 | 2.9 | 70 |
| ATLP1-S37 | 17.1 | 35.7 | 18.6 | 21.4 | 7.1 | 70 |
| ATLP1-As8 | 16.9 | 22.5 | 11.3 | 19.7 | 29.6 | 71 |

<sup>a</sup>Sporulation classes are as follows: 0= no infection/ non-sporulating lesions, 1= 1-25% lesion covered with spores, 2= 26-50% lesion covered with spores, 3= 51-75% lesion covered with spores, 76-100% lesion covered with spores.
ATLP1-S3 to S37 are sense transgenic lines whereas, ATLP1-As8 is a control line

**Figure 3.**
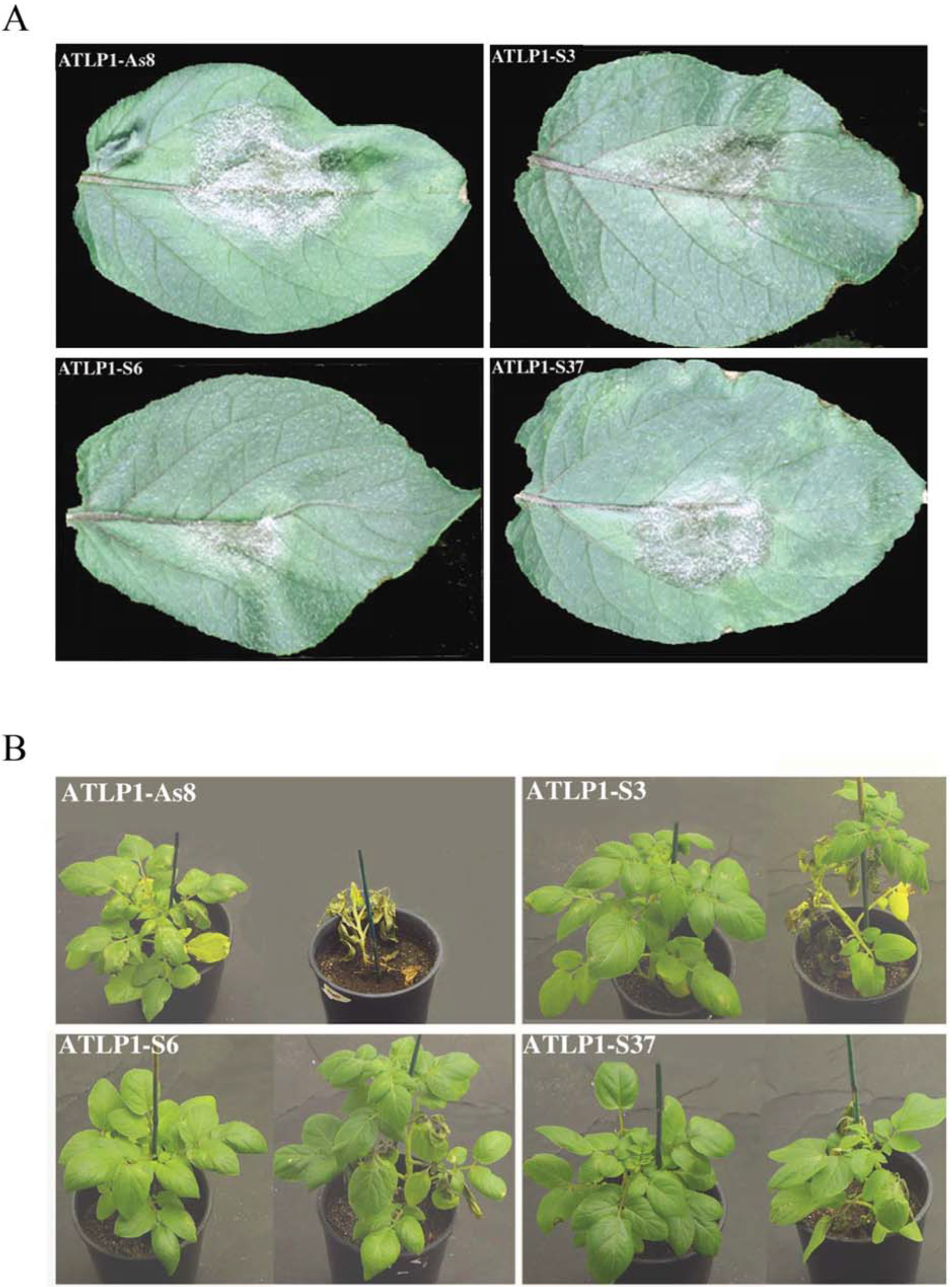
Resistance of ATLP1-transgenic potato plants to *P*. *infestans*. **A)** Detached leaflets were inoculated with sporangial suspension of *P*. *infestans* and placed at 17°C. ATLP1-S3, S6 and -S37 are ATLP1-transgenic lines; ATLP1-As8 is a control line. Inoculated leaflets were scanned and disease area measured. Photographs given are representative from two independent experiments. **B)** Resistance of ATLP1-transgenic potato plants to *P*. *infestans*. Whole plants were spray-inoculated with a sporangial suspension of *P*. *infestans*, and placed at 17°C. Pictures on the left and right in each panel were taken 10 and 20 days post-inoculation, respectively. Pictures are representative of two independent experiments consisting of five to eight pots per line.

In whole plant assays four ATLP1-transgenic lines showed enhanced resistance against *P*. *infestans*. Symptoms usually appeared 4 days after inoculation and continued to expand. However, no difference between control and ATLP1-transgenic lines could be detected at this stage. Ten days after inoculation numerous water soaked lesions were observed on control plants as compared to fewer and smaller lesions on ATLP1-transgenic plants (Figure 3B, left plant in each panel). The percent infected leaf area of each ATLP1-transgenic line was compared to control using the student t-test. Four lines had significantly lower disease than control (Table 4). The level of disease reduction ranged from 40 to 70% (Table 4). The disease lesions continued to expand and coalesce. Twenty days after inoculation most of the control plants were severly infected with dead leaves whereas ATLP1-transgenic lines had fewer dead leaves and most of the plants remained alive with reduced disease level (Figure 4B, right plant in each panel).

**Table 4.** Resistance of ATLP1-trasgenic potato plants to *P*. *infestans*.

| **Transgenic** | **% reduction** |  |
| --- | --- | --- |
| **Lines** | **over control** | **p-value<sup>a</sup>** |
| ATLP1-S3 | 67.7 | 0.0286 |
| ATLP1-S6 | 70.0 | 0.0007 |
| ATLP1-S11 | 40.8 | 0.2629 |
| ATLP1-S13 | 45.4 | 0.0476 |
| ATLP1-S37 | 65.4 | 0.0727 |
| ATLP1-As8 | 00.0 | - |

<sup>a</sup> The differences between ATLP1-As8 and ATLP1-S lines were tested by student's T-test. P-values are probabilities of getting a larger value of T if the difference between an ATLP1-S transgenic and ATLP1-As8 control was truly equal to zero

## DISCUSSION

The Arabidopsis thaumatin-like protein 1 (ATLP1) was selected for genetic improvement of potato, because it is induced in response to pathogen attack and by compounds that are known to elicit systemic acquired resistance. The ATLP1 gene used in this study was cloned from a cDNA library of Arabidopsis, and characterized to belong to the PR5 family of pathogenesis related proteins (Hu and Reddy, 1997). PR5 proteins, which share considerable sequence similarities to the extremely sweet protein, thaumatin, from *Thaumatococcus daniellii* fruits (Cornelissen et al., 1986), have been shown to have antifungal activities e.g. against *P*. *infestans*, both *in vitro* (Woloshuk et al., 1991) as well as when expressed in transgenic plants (Liu et al., 1994).

In this study the ATLP1 gene under the 35S promoter has been introduced into potato Cv. Desiree. Over 50 independent transformants were obtained. Several of these had substantially higher levels of expression of the transgene at the mRNA and protein levels. The ATLP1 expression was different in different independent lines, which is consistent with the previously reported variable expression of a rice thaumatin-like protein when expressed in rice and wheat (Chen et al., 1999; Datta et al., 1999). This could be due to the positional effect of the introduced gene. The high expresser lines were selected for response to inoculation with *A*. *solani* and *P*. *infestans*-the two most damaging fungal pathogens of potato worldwide (Hooker, 1981).

Most of the previous studies with transgenic plants that have reported enhanced disease resistance have employed chitinase genes (Broglie et al., 1991; Howie et al., 1994; Zhu et al., 1994; Jach et al., 1995; Punja and Raharjo, 1996). Since chitinases are effective against only those fungal pathogens that contain chitin in their cell wall, they can not be used more effectively against *P*. *infestans*, which contain very little or no chitin (Fry and Goodman, 1997). The exact mechanism of the inhibitory effect of ATLP1 is not known completely. However, studies with proteins that share substantial sequence similarity to ATLP1, indicate that permeability of cell membranes is compromised (Roberts and Selitrennikoff, 1990; Vigers et al., 1991; Abad et al., 1996). Alternatively, ATLP1 might be acting in a two-step mechanism as proposed for several thaumatin like proteins (Trudel et al., 1998). According to this mechanism, thaumatin–like proteins could bind to β-1,3 glucans of fungal wall as a prerequisite for subsequent insertion into cell membranes to cause trasmembranes pores (Roberts and Selitrennikoff, 1990; Trudel et al., 1998). A third proposed mechanism of the action could be the direct enzymatic degradation of polymeric ß-1,3 glucans, found in fungal cell walls (Grenier et al., 1999). Nevertheless, the availability of a gene with a mode of action different than chitinases should prove useful for producing transgenic potato plants resistant to fungal pathogens.

Potato plants, like other plants, should posses the ability to mount a defense reaction when challenged by pathogens. However, in susceptible plants these defense mechanism are suggested to be too weak and/or slow to provide timely and effective protection against the invading pathogens (Fritig et al., 1998). Moreover, due to the continuous co-existence of host and pathogens, a pathogen could evolve to counteract or avoid the toxic effects of plant antifungal compounds (Lamb et al., 1992). Under these circumstances, antifungal products from a taxonomically distant plant species, which is not a host for a pathogen, might prove more effective. Since, the transfer of antifungal genes from distantly related plant species is often not possible by conventional breeding methods, genetic engineering is probably the best strategy to accomplish the transfer of such genes to crops. ATLP1, used in this study, has been cloned from Arabidopsis, which is not normally infected by *P*. *infestans* or *A*. *solani*. Therefore, it was proposed that the expression of ATLP1 in potato could enhance resistance against these fungal pathogens. In order to test this hypothesis, we introduced ATLP1 in potato plants, which showed enhanced fungal resistance.

The transgenic plants showed different degree of protection against *A*. *solani* and *P*. *infestans*. Symptom development for early blight and late blight were reduced roughly by 53-82%, and 4-24% (detached-leaf assay) and 40-70% (whole-plant assay), respectively. These different degrees of protection might be due to differences in the susceptibility of these two organisms to ATLP1. A rice thaumatin-like protein when expressed in rice and wheat also gave different level of protection against *Rhizoctonia Solani* and *Fusarium graminearum* (Chen et al., 1999; Datta et al., 1999). Also, the variability observed in the expression of protein and the level of disease resistance of independent ATLP1-transgenic potato lines is in conformity with the results reported with transgenic rice and wheat expressing a rice thaumatin-like protein (Chen et al., 1999; Datta et al., 1999). As demonstrated by reduction in IE, LS and SC in transgenic potato plants, the mechanism of protection seems to affect different stages in the life cycle of *P*. *infestans*. Infection efficiency, symptom development and sporulation capacity are the parameters that determine rate and size of epiphytotic developments (Parlevliet, 1979). Their reduction in the ATLP1 transgenic plants could result in reduced blight diseases of potato. Under greenhouse conditions no undesirable phenotypic characteristics were observed in ATLP1-transgenic lines.

Disease response in transgenic plants is influenced by the manner in which plant tissues are colonized by the invading fungus and by the location of the antifungal protein. For example in tobacco, overexpression of a vacuolar chitinase had no effect on *Cercospora nicotianae* (Neuhaus et al., 1991; Nielsen et al., 1993), a pathogen that grows intercellularly and would not encounter the intracellular chitinase. Similarly, a biotrophic or hemibiotrophic pathogens that grow intercellularly might not be expected to be affected by a product accumulating in vacuoles. The ATLP1-protein, because of the presence of a secretory amino-terminal signal peptide, is secreted to the extracelullar spaces in potato plants (Figure 1C). In this study, *P*. *infestans*, which is a hemibiotrophic oomycete, and *A*. *solani*, which is necrotrophic and grows intercellularly, are shown to be inhibited by the ATLP1 most probably accumulated in the intercellular spaces of transgenic potato plants. These results demonstrate that the Arabidopsis thaumatin-like protein 1 may prove effective against pathogens from distantly-related taxonomic groups.

As shown here in this study and others reported elsewhere, complete protection by the introduction of a single defense related gene is not achieved. The expression of resveratrol synthase in tomato (Thomzik et al., 1997) and of osmotin-like protein in potato (Liu et al., 1994; Zhu et al., 1996) resulted in enhanced resistance against *P*. *infestans*. Since coexpression of two antifungal genes are reported to confer higher resistance than either gene alone (Zhu et al., 1994; Jach et al., 1995), the expression of a combination of genes, preferably with different mode of actions e.g. ATLP1 (this study), ribosome inactivating protein (Jach et al., 1995), osmotin-like protein (Zhu et al., 1996) and resveratrol synthase (Thomzik et al., 1997) or synthetic antimicrobial peptides (Ali and Reddy, 2000; Ali et al., 2003) will provide greater and broad-spectrum protection against fungal pathogens in potato.

## Authors contribution

AR conceived and designed research. GA and XH conducted experiments. AR, GA and XH analyzed data and wrote the manuscript. All authors read and approved the manuscript.

## Acknowledgments

We are thankful to Dr. Joe Hill and Dr. Howard Shwartz of Colorado State University, Fort Collins, for providing the *Alternaria solani* and *Phytophthora infestans* strains; Dr. An for the pGA748 vector. This work was supported by the Colorado Institute for Research in Biotechnology, Colorado Agricultural Experiment Station and Colorado Potato Administrative Committee (Area II).

